# Improve the Efficacy of B7-H3-Targeting Antibody-Drug Conjugate DS-7300a in TP53-deficient Tumors by Inducing Ferroptosis

**DOI:** 10.64898/2026.05.05.722958

**Authors:** Javier Leo, Feiyu Chen, Wei Shi, Xin Liang, Chenling Meng, Qianlin Gu, Yaman Albittar, Zhen Fan, Jie Zhang, Boyi Gan, Sangeeta Goswami, Kendra Carmon, Daniel E. Frigo, Ana Aparicio, Di Zhao

**Author notes:** **Corresponding Author**: Di Zhao, Department of Experimental Radiation Oncology, The University of Texas MD Anderson Cancer Center, 6565 MD Anderson Boulevard, Houston, TX 77030, USA.

## Abstract

Immune checkpoint B7-H3 is an emerging target for immunotherapy. DS-7300a is an advanced B7-H3-targeting antibody-drug conjugate (ADC) warheaded with the topoisomerase I inhibitor DXd. DS-7300a has demonstrated clinical activity, but molecular biomarkers to predict its therapeutic response remain elusive. *TP53* is one of the most mutated tumor suppressor genes across cancers, and effective therapies are urgently needed for *TP53*-deficient cancers. Using prostate cancer (PCa) as a model system, we reported that DS-7300a’s anti-tumor efficacy is highly dependent on functional p53 in cancer cells, and *TP53* defects confer resistance to DS-7300a. Mechanistically, we found that DS-7300a and its payload, DXd, induce DNA damage and activate the ATM/ATR/CHK signaling cascade, thereby stabilizing p53 and inducing a pro-apoptotic and senescence-associated transcriptome. In contrast, *TP53*-deficient cells fail to detect DXd-induced DNA damage, maintain a high proliferation rate, and exhibit low levels of apoptosis and senescence, thereby conferring resistance to DS-7300a. Ferroptosis is an iron-dependent form of regulated cell death triggered by lipid peroxidation, which is mechanistically and morphologically distinct from apoptosis. Interestingly, DS-7300a treatment elevates lipid peroxidation in TP53-deficient cancer cells and upregulates glutathione peroxidase 4 (GPX4), an antioxidant enzyme that mitigates lipid peroxidation. Using isogeneic xenograft models and a newly developed humanized B7-H3 PCa model, we demonstrated that inducing ferroptosis by pharmacological inhibition of GPX4 enhances DS-7300a’s efficacy in *TP53*-deficient tumors. Our studies demonstrate that TP53 status dictates anti-tumor responses to DS-7300a, and ferroptosis induction represents a promising therapeutic approach to overcome resistance to DS-7300a in malignancies harboring TP53 defects.

## Introduction

The tumor suppressor TP53 plays a central role in regulating cell division, cell death, and the response to DNA damage [1–3]. Genetic mutations or deletions in *TP53* are found in more than half of cancers [4–8] and are associated with higher metastatic rates and poorer clinical outcomes. Prostate cancer (PCa) is the most diagnosed cancer and the second leading cause of cancer death in men in the United States [9]. While most patients initially respond to androgen deprivation therapy, their disease often progresses to castration-resistant prostate cancer (CRPC) [10, 11]. Together with PTEN and RB1, *TP53* defects have been used to characterize a clinical-biological subset of androgen-indifferent prostate tumors, termed aggressive variant prostate cancers (AVPC) [8]. Treatment options for advanced PCa with TP53 defects are limited. Effective therapeutics and combinatorial strategies are urgently needed.

Antibody-drug conjugates (ADCs) comprise antibodies conjugated to cytotoxic payloads via a cleavable linker. After the ADC binds to the cell-surface antigen, the ADC-antigen complex undergoes internalization and translocation to the lysosome, where the ADC is cleaved, releasing the payload drug, resulting in cancer cell death [12]. ADCs are designed to exert anti-tumor activity in a target-dependent manner and reduce systemic toxicity. DNA topoisomerase 1 (TOP1) plays an essential role in DNA replication, transcription, recombination, and chromosome condensation by relaxing DNA supercoiling. Inhibitors targeting TOP1 bind to the DNA-TOP1 complex and lead to DNA strand breaks and cell death [13]. The exatecan derivative (DXd) is a novel, selective TOP1 inhibitor payload that has been used to develop several ADCs currently under clinical investigation [14–18]. Among them, the HER2-targeting ADC trastuzumab deruxtecan demonstrated robust efficacy in preclinical and clinical studies [14, 19–21]. It has recently been approved in many countries, including the United States, for treating multiple HER2-positive breast cancers.

Immune checkpoint B7-H3 (*CD276* gene) is a member of the B7 and CD28 families. B7-H3 overexpression is correlated with increased risks of disease progression and poor outcomes in cancer patients [22–29]. Several therapeutic approaches to targeting B7-H3 have been tested in preclinical and clinical studies, including monoclonal antibodies (mAbs), ADCs, and chimeric antigen receptor T cells (CAR-T) [28, 30–35]. Our prior studies demonstrate that high B7-H3 expression in cancer cells is induced by TP53/PTEN defects and contributes to immunosuppression and prostate tumor progression [36], underscoring B7-H3 as a promising therapeutic target in cancers containing TP53 defects.

DS-7300a is among the most advanced B7-H3-targeting ADCs. DS-7300a comprises a humanized anti-B7-H3 IgG1 mAb warheaded with TOP1 inhibitor DXd (**Fig. 1A**) [16, 37]. DS-7300a exhibited potent anti-tumor effects in preclinical models [16, 37] and showed promising clinical activity in patients with metastatic CRPC, small cell lung cancer, and other malignancies (NCT04145622 and NCT05280470). However, the molecular subsets of cancer patients that benefit most from B7-H3-targeting ADC remain unknown. Prior studies have shown that the anti-tumor efficacy of DS-7300a correlates positively with higher B7-H3 expression in cancer cells [16]. Given that TP53 defects induce B7-H3 expression, we hypothesized that TP53-deficient tumors are more sensitive to DS-7300a.

**Figure 1.**
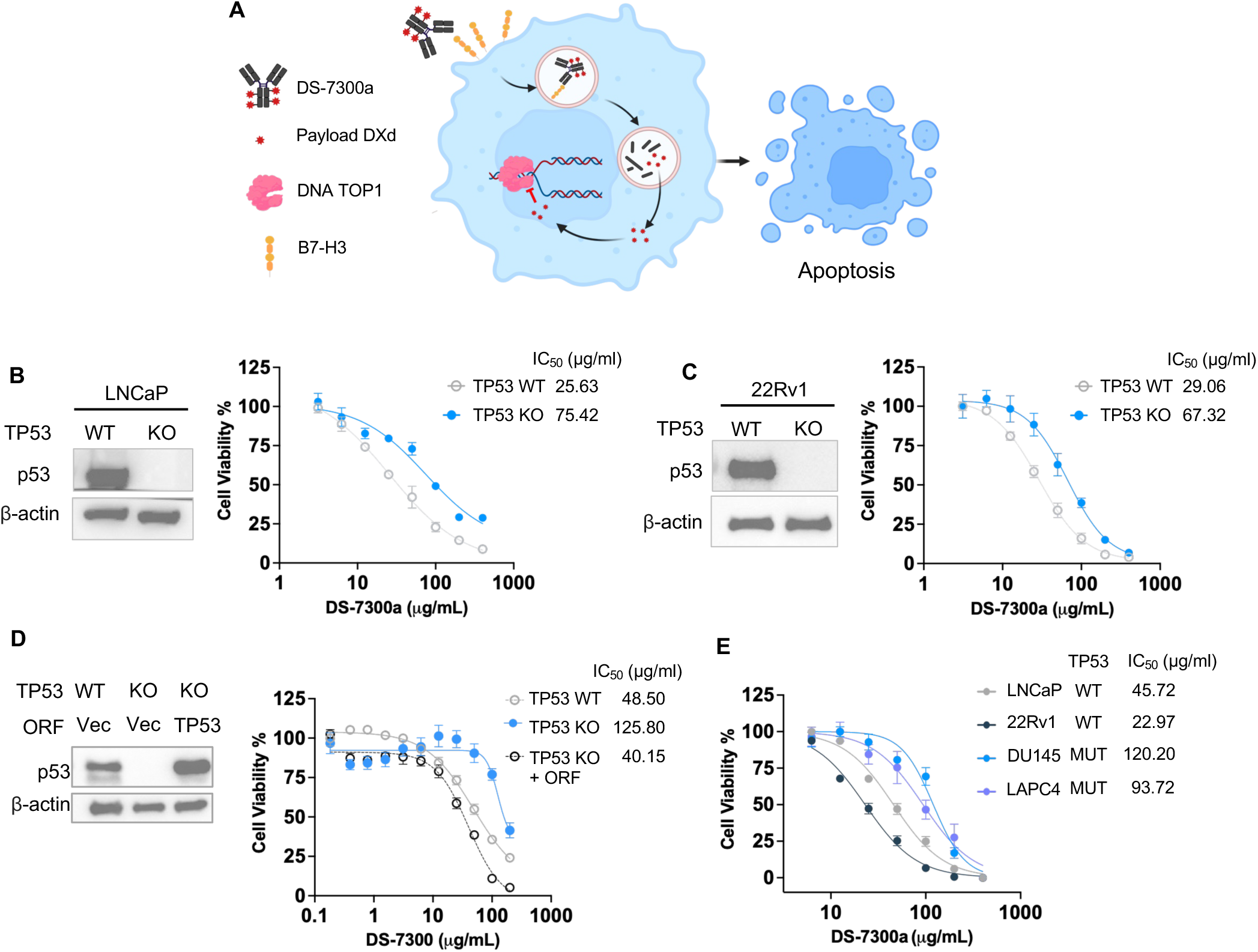
TP53 defects confer resistance to DS-7300a, a B7-H3-targeting ADC. **A.** Schematic of the mechanism of action of B7-H3-targeted antibody-drug conjugate, DS-7300a equipped with a DNA topoisomerase I (TOP1) inhibitor, Dxd. **B-C.** Western blot of p53 and dose-response curves of DS-7300a in TP53-WT versus TP53-KO LNCaP (B) and 22Rv1 (C) cells. **D.** TP53 was reintroduced into TP53-KO LNCaP cells, followed by Western blot and DS-7300a dose-response assays. **E.** IC_50_ determination of DS-7300a in diverse PCa cell lines containing wildtype or mutated *TP53*. Cells were treated with the indicated concentrations of DS-7300a for 6 days in all experiments with at least three replicates. Data represent the mean ± standard deviation of triplicates or otherwise stated. IC_50_ values were calculated using GraphPad Prism version 10.4.1.

In this study, we used PCa as a model system to evaluate the activity of B7-H3-targeting ADC DS-7300a in tumors with or without TP53 defects. Surprisingly, we found that TP53 defects rendered cancer cells more resistant to DS-7300a *in vitro* and *in vivo*. Moreover, we uncovered the mechanisms underlying this resistance and developed a novel combination with ferroptosis inducers to improve the anti-tumor efficacy of DS-7300a in cancers with TP53 deficiency.

## METHODS

### Study Design

This study determined the efficacy of the B7-H3 ADC, DS-7300a, understanding the mechanisms of resistance, and provided new insights into combination therapies to overcome this resistance. To this end, we tested DS-7300a and its payload, DXd, in various cell lines containing varying p53 statues and performed *in vitro* assays (Western blot, quantitative real-time PCR, cell viability assays, and IHC staining), and *in vivo* experiments using syngeneic xenograft and humanized models. For the *in vitro* cell assays, data were collected from at least three independent cultures, as indicated in the figure legends. Due to the nature of PCa, only male mice were used for *in vivo* experiments, with biological replicates indicated in figure legends. Sample sizes were determined based on previous experience for each experiment. Mice were randomly assigned to experimental groups whenever possible. Tumor progression was evaluated by tumor size throughout the course of treatment and by tumor weight at the endpoint. Tumor-bearing mice were euthanized when the tumor volumes reached approximately 2000 mm³. Analysis of each experiment was done blindly when possible. All mouse experimental procedures followed the Institutional Animal Care and Use Committee (IACUC) protocol (#00001955). The mice were housed under a 12-hour light/dark cycle, at 68-78°F with 30–70% humidity. MD Anderson IACUC’s guidelines for the proper and humane use of animals in biomedical research were followed. Euthanasia was carried out at the endpoint by CO_2_ inhalation followed by cervical dislocation, which is an approved method of euthanasia for mice.

### Cell Culture

HEK293T and DX1 cells were cultured in DMEM medium (Corning, 10-013CV) with 10% FBS (Thermo Scientific, 10082147) and 1% penicillin-streptomycin (Thermo Scientific, 15140163). 22RV1 and LNCaP cells were cultured in RPMI-1640 medium (Corning, 10-040-CV) with 10% FBS (Thermo Scientific, 10082147) and 1% penicillin-streptomycin (Thermo Scientific, 15140163). DU145 was cultured in EMEM medium (ATCC, 30-2003) with 10% FBS (Thermo Scientific, 10082147) and 1% penicillin-streptomycin (Thermo Scientific, 15140163). Cell lines were authenticated by MD Anderson’s Cytogenetic and Cell Authentication Core and were regularly tested for mycoplasma using MycoAlert PLUS detection kit (Lonza, LT07-710), according to the manufacturer’s instructions.

### Transient and Lentiviral Transfection

Transient transfection of siRNA or plasmids (Supplementary Table 1) was performed using Lipofectamine 2000 transfection reagent (Thermo Fisher Scientific, 11668019) according to the manufacturer’s instructions. After 48 hours, the cells were used for further analyses. Human p53 overexpression plasmid (Horizon Discovery, OHS5893-202493665) was transfected into p53-KO LNCaP cells using Lipofectamine 2000. Lentivirus was used to establish stable cell lines for CRISPR-mediated *CD276* knockout or gene overexpression. Empty lentiviral vectors were used as the control. For lentiviral transduction, lentiviral plasmids were transfected into 293T cells together with lentiviral expression constructs, psPAX2, and pMD2.G. After 48 h, the viral supernatants were harvested and filtered and then used to infect DX-1 cells with 10 μg/mL polybrene (EMD Millipore, TR-1003-G) and subsequently selected with 10 μg/mL puromycin (Gibco, A1113803) transduction reagent.

### In Vitro Inhibitor Treatment

TP53-KO LNCaP cells were treated with DXd (DMSO, 20nM and 40nM) for 48 hours or DS-7300a (ABS, 100ug/mL, 200ug/mL) for 120 hours, followed by determination of protein or gene expression using Western blot analysis and qPCR, respectively.

### Cell Viability Assays and IC50 Determination

Respective PCa cell lines were seeded into 96-well plates and then treated with DXd or DS-7300a at serially diluted concentrations. For DS-7300a, cells were treated at maximal concentrations of 400ug/mL or 40nM for DXd, then serially diluted by a factor of two, unless otherwise noted in figure legends. For IC50 determination, the CellTiter-Glo Luminescent Cell Viability Assay kit (Promega, G7573) was used, and the viability of cells in each well was standardized against the DMSO control group. The IC50 values were analyzed using GraphPad Prism version 9.2.0. The synergistic effects between DS-7300a and RSL-3 were determined using CellTiter-Glo and analyzed using SynergyFinder (v3.0). Bliss Synergy Scores of the drug combination were analyzed.

### Western Blot Analysis

The cell pellets were then lysed in 1x Laemmli sample buffer (Bio-Rad Laboratories, 1610747), premixed with 2-mercaptoethanol (Bio-Rad Laboratories, 1610710), at 99 °C for 20 min. Proteins were separated with 4%−15% Mini-PROTEAN TGX Precast Protein Gels (BioRad, 4561086) and transferred to the nitrocellulose membrane using the Trans-Blot Turbo RTA Mini 0.2-µm nitrocellulose transfer kit (Bio-Rad Laboratories, 1704270). The blots were then blocked in 0.5% milk in TBST buffer containing 0.01% Tween-20, then incubated with primary antibodies overnight at 4 °C. The primary antibodies employed are as follows: anti-B7-H3 (R&D Systems AF1027, 1:1000), anti-TOP1 (Cell Signaling Technologies, 79971, 1:1000), anti-phospho-H2Ax (Cell Signaling Technologies, 2577, 1:1000), anti-phospho-CHK2 (Cell Signaling Technologies, 82263T, 1:1000), anti-p53 (Santa Cruz, sc-126, 1:1000), anti-p21 (Cell Signaling Technologies, 37543, 1:1000), anti-GPX4 (R&D Systems, MAB5457, 1:500) and anti-β-Actin (Sigma-Aldrich, A5441, 1:10000) (**Supplementary Table 1**). Respective HRP-conjugated secondary antibodies were diluted in 0.5% milk in TBST and incubated with the membranes for 1 hour. Colorimetric and chemiluminescence signals were developed by Western ECL substrates (Bio-Rad Laboratories, 1705060) and captured using the ChemiDoc Imaging System (Bio-Rad Laboratories).

### Quantitative Real-time PCR

Total RNA was isolated from cell samples using the RNeasy Mini Kit (Qiagen, Inc., 74106). Complementary DNA (cDNA) was then synthesized via reverse transcription using the High-Capacity cDNA Reverse Transcription Kit (Life Technologies, 4368813), following the manufacturer’s standard protocol. Quantitative real-time PCR (qPCR) was performed on a QS3 Real-Time PCR System (Thermo Fisher Scientific) with PowerUp SYBR Green Master Mix (Thermo Fisher Scientific, A25778), in accordance with the manufacturer’s instructions. Gene relative expression was quantitatively determined using indicated primers (**Supplementary Table 1**). The relative mRNA expression levels were normalized to actin controls. All the data were analyzed using GraphPad Prism version 10.4.1.

### Lipid Peroxidation Assay

Cells were seeded in 24-well plates and treated with DMSO or DS-7300a. After 72hr, cells were collected by trypsinization and resuspended in 500 μL PBS containing 5 μM C11-BODIPY 581/591 (Invitrogen, D3861). Following a 30-minute incubation at 37°C, lipid peroxidation percentages were quantified by flow cytometry using oxidized C11-BODIPY 581/591 fluorescence (excitation: 488 nm; emission: 530/30 nm).

### β-galactosidase Assay

LNCaP cells were seeded in 6 or 24-well plates and treated with DXd (0 and 40nM) or DS-7300a (0 and 200 μg/mL) for 24 and 108 hrs, respectively. Cells were then washed and stained using a β-galactosidase staining kit (Cell Signaling Technology, 9860S). Cells were imaged using the Cytation 5 imager (BioTek), and positive cells were counted using QuPATH (v0.6.0-rc5) in quadruplicate views.

### Tumor Implantation and Treatment in Xenograft Models

2×10^6^ LNCaP cells with or without *TP53* knockouts [36] were subcutaneously injected into both flanks of seven-week-old male SCID mice (CB17 Charles River). When tumors reached approximately 50-70mm^3^, tumor-bearing mice were randomized into groups to receive vehicle or 0.5 mg/kg DS-7300a via tail vein injection. Mice were treated with DS-7300a biweekly, twice. Tumor size was measured every three days using calipers. Tumor volume was calculated (width × width × length/2). At the endpoint, tumors were collected for histopathologic analyses. To test the combination of DS-7300a and JKE-1674 in TP53-deficient tumors, 2×10^6^ TP53-KO LNCaP cells were subcutaneously injected into both flanks of seven-week-old male SCID mice. When tumors reached approximately 50-70mm^3^, tumor-bearing mice were randomized into two groups for combination treatment (3 mg/kg DS-7300a, i.v., biweekly, twice; 15 mg/kg JKE-1674, oral gavage, every two days) or vehicle control. Tumor growth was monitored, and tumor weight was measured.

### Generation of a new humanized B7-H3 PCa model

B-hB7-H3 mouse strain with C57BL/6J background was purchased from Biocytogen (Cat #110028). The mice were bred and housed at MD Anderson, where they were monitored daily for health. Regular genotyping was conducted for all GEMM mice throughout the study. The human B7-H3 gene was introduced into the syngeneic DX1 PCa cell line to generate DX1-hCD276 cells, and B7-H3 protein expression was assessed by Western blot and Flow Cytometry. 2×10^6^ DX1-hCD276 cells were subcutaneously injected into both flanks of male B-hB7-H3 mice with 1:1 Matrigel. When tumor sizes reached 50-80 mm^3^, tumor-bearing mice were randomly assigned for treatment with ABS vehicle control or DS-7300a at two doses (0.5 mg/kg or 3 mg/kg; i.v., biweekly, twice). The body weights of the mice were measured weekly. Tumor growth was measured every three days using calipers. Tumor volume was calculated (width × width × length/2). For combination treatment, tumor-bearing mice were randomly assigned for single or combination treatment of DS-7300a (2 mg/kg; i.v.; once) or/and JKE-1674 (10 mg/kg; oral gavage, every two days). Mice from each cohort were euthanized at the endpoints to collect tumor samples, followed by histopathology analysis.

### Immunohistochemistry (IHC)

The tissues or tumors were dissected and embedded in paraffin for H&E and immunohistochemistry staining. Paraffin-embedded slides were incubated at 60°C and then rehydrated. Antigen retrieval was performed with citrate-based antigen unmasking solution (Vector Laboratories, H-3300-250) at 95°C for 30 min and cooled down to room temperature. Sections were blocked with 10% normal donkey serum (NDS, Sigma-Aldrich, 566460-5ML) for 1 hour. Primary antibodies were prepared in antibody diluent (Agilent Technologies, S080983-2) overnight at 4°C. The primary antibodies employed are as follows: anti-B7-H3 (R&D Systems AF1027, 1:1000), anti-phospho-H2Ax (Cell Signaling Technologies, 2577, 1:1000), anti-p53 (Santa Cruz, sc-126, 1:1000), anti-p21 (Cell Signaling Technologies, 37543, 1:1000), anti-GPX4 (R&D Systems, MAB5457, 1:500) and anti-Ki67 (ThermoFisher Scientific, RM-9106-S1 1:5000) (**Supplementary Table 1**). The slides were scanned by Aperio CS2 Scanscope (Leica). At least five views (20× magnification) per tumor were captured in a blinded manner using QuPATH (v0.6.0-rc5). Quantification plots represent the mean ± standard deviation of >5 individual views per tissue section. Statistical analyses were performed using GraphPad Prism version 10.4.1.

### Statistical analyses

The number of samples is shown in figures and described in figure legends. All data are presented as the mean ± standard deviation of at least triplicate experiments, unless otherwise stated. No statistical method was used to determine the sample size in advance. Comparisons between the two groups were performed using an unpaired, two-tailed Student t-test. One-way analysis of variance (ANOVA) with Tukey’s post hoc tests was performed to analyze data from three or more groups, as indicated in the figure legends. Statistical analysis was performed using GraphPad Prism version 10.4.1. Statistical comparisons between the two groups were conducted using the unpaired two-tailed Student’s t-test. For comparisons with more than two groups, one-way ANOVA followed by Tukey’s post hoc test was applied, as described in the figure legends. For all experiments, statistical significance is defined as follows: ****P < 0.0001, *** P < 0.001, ** P < 0.01, * P < 0.05. ns, not significant.

## RESULTS

### *TP53* defects confer resistance to DS-7300a, a B7-H3-targeting ADC

To determine the activities of DS-7300a, we first generated *CD276* knockout isogenic PCa cells using the sgRNA-guided CRISPR/Cas9 system and determined the dose response of DS-7300a. As shown in **Fig.S1A**, the depletion of B7-H3 reduced the cytotoxic activity of DS-7300a in LNCaP cells, indicating the on-target effect of the ADC drug. To assess the impact of TP53 on DS-7300a efficacy, we genetically knocked out *TP53* in LNCaP and 22Rv1 cells that contain wild-type TP53. TP53 knockout cells exhibited a 2-3-fold increase in IC_50_ values compared with wild-type control cells, indicating reduced sensitivity to DS-7300a (**Fig. 1B, C**). This reduced sensitivity was reversed entirely by re-introducing TP53 overexpression construct, underscoring the key role of p53 in mediating the cytotoxic activity of DS-7300a (**Fig. 1D**). To evaluate the associations between TP53 status and the response to DS-7300a, we determined the IC_50_ of DS-7300a in a panel of PCa cells containing wild-type TP53 (LNCaP and 22Rv1) or mutated TP53 (DU145 and LAPC4). As shown in **Fig. 1E**, TP53-deficient cancer cells presented a 3-4-fold increase in IC_50_ values, compared to TP53-intact cells. Of note, DS-7300a’s sensitivity was associated with TP53 status rather than B7-H3 expression (**Fig. S1B**).

Next, we determined the effects of DS-7300a in xenograft PCa models derived from TP53-wildtype and TP53-knockout LNCaP cells (**Fig. 2A**). The results showed that DS-7300a effectively suppressed the growth of TP53-wildtype tumors but had modest effects on TP53-knockout tumors (**Fig. 2B-C and S1C**). Immunohistochemistry analysis revealed that DS-7300a treatment reduced cancer cell proliferation in TP53-intact tumors, as featured by Ki67, but not in TP53-deficient tumors (**Fig. 2D**). Given the immunomodulatory role of B7-H3, we sought to assess the anti-tumor activity of DS-7300a in an immunocompetent PCa model. Due to DS-7300a’s specificity to human B7-H3, we generated a novel humanized B7-H3 PCa mouse model (**Fig. 2E**). Briefly, we introduced the human B7-H3 into our well-characterized syngeneic PCa cell line DX1 containing co-deletion of *Pten/Trp53/Smad4* and generated a hB7-H3-DX1 cell line expressing membrane-bound human B7-H3 protein (**Fig. 2F**). Then, we injected these hB7-H3-DX1 cells into the humanized B7-H3 mice with a C57BL/6 background (**Fig. 2E**). After the tumor sizes reached 50 mm^3^, we treated the mice with low dose (0.5 mg/kg) or high dose (3 mg/kg) of DS-7300a. The results indicated that TP53-deficient hB7-H3-DX1 tumors were resistant to DS-7300a treatment (**Fig. 2G-H and S1D-E**). The ADC drug displayed little toxicity *in vivo* at both doses (**Fig. S1F**). Histopathology analysis showed that DS-7300a reduced the expression of the target protein B7-H3 and induced DNA damage, as featured by yH2Ax, but failed to reduce cell proliferation in TP53-deficient tumors (**Fig. S1G**). Taken together, these *in vitro* and *in vivo* studies across various PCa cell lines and models demonstrate that TP53 defects render cancer cells more resistant to DS-7300a.

**Figure 2.**
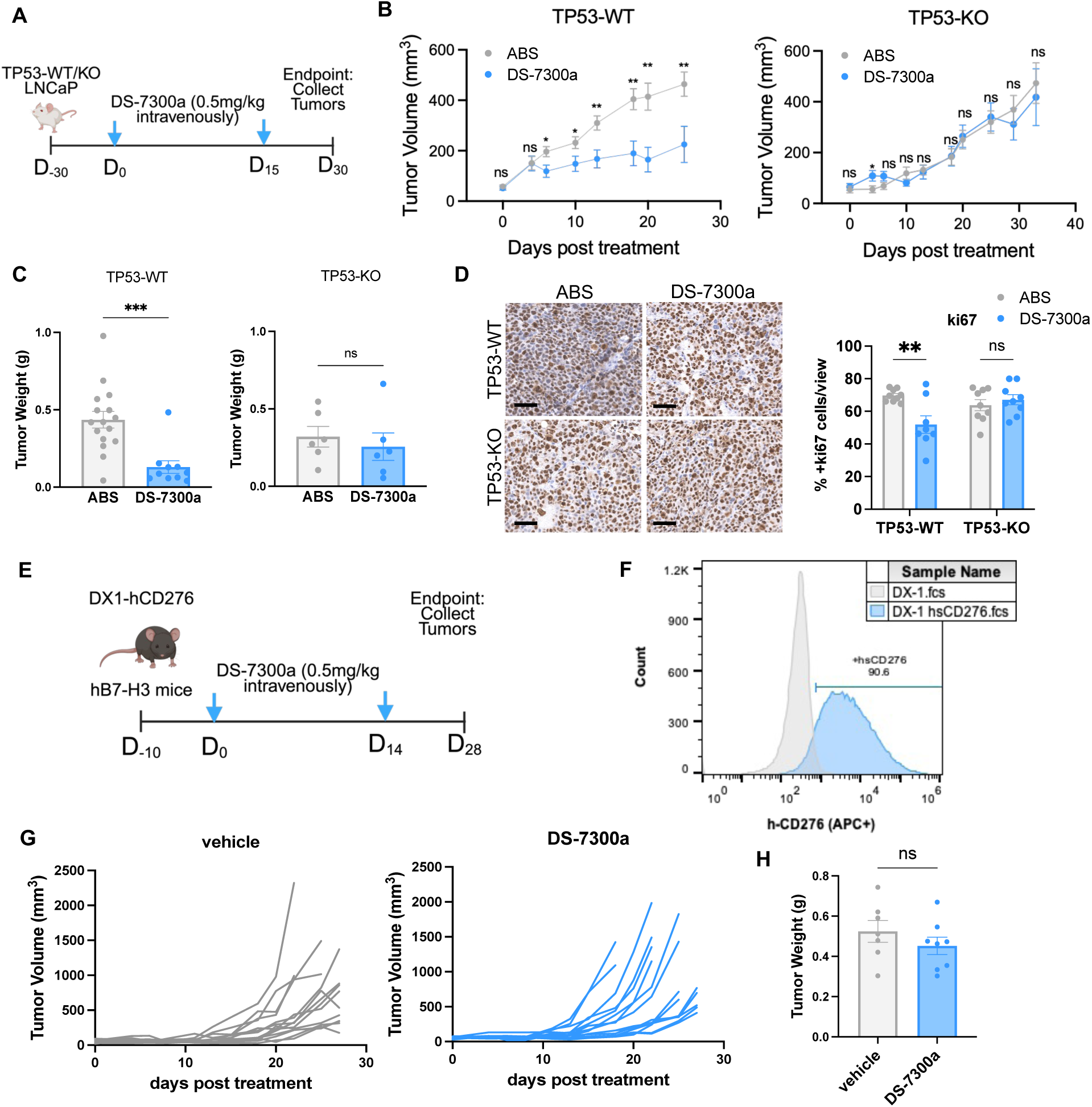
TP53-deficient tumors showed resistance to DS-7300a in xenograft and humanized models. **A.** Schematic of experimental design using xenograft models. 2×10^6^ TP53-WT and TP53-KO LNCaP cells were subcutaneously injected into both flanks of seven-week-old male SCID mice. When tumors reached approximately 75mm^3^, tumor-bearing mice were randomized for treatment of ABS vehicle or DS-7300a (0.5 mg/kg; i.v., biweekly, twice). **B-C.** Tumor growth over time (B) and tumor weights at the endpoint (C) in xenograft models are shown. **D.** Representative images and quantification of Ki67 IHC staining in xenograft tumors after treatments. Scale bar = 50um. **E.** Schematic of experimental design using humanized B7-H3 models. Human B7-H3 gene was introduced into the syngeneic DX1 PCa cell line to generate DX1-hCD276 cells. 2×10^6^ DX1-hCD276 cells were subcutaneously injected into both flanks of male B-hB7-H3 mice, followed by treatment of ABS vehicle control or DS-7300a (0.5 mg/kg; i.v., biweekly, twice). **F.** B7-H3 protein expression of DX1-hCD276 cells was verified by Flow Cytometry. **G-H.** Tumor growth over time (G) and tumor weights at the endpoint (H) of humanized B7-H3 PCa models after treatment. Data represent the mean ± standard deviation of triplicates or otherwise stated. Statistics were calculated using GraphPad Prism version 10.4.1. *P* values determined by unpaired two-tailed t test (B, C, H) or by two-way ANOVA analysis (H); *P < 0.05; **P < 0.01; ***P < 0.001; ****P < 0.0001; ns, not significant.

### DS-7300a’s payload DXd triggers apoptosis and senescence via activating p53

Furthermore, we investigated whether DS-7300a resistance was attributable to the target B7-H3 or to the DNA TOP1 inhibitor payload, DXd. Phenocopying DS-7300a, DXd exhibited higher IC50 values in TP53-deficient LNCaP cells (**Fig. 3A**) and 22Rv1 cells (**Fig. S2A);** but this effect was abrogated with the reintroduction of TP53 (**Fig. 3B**). The associations between TP53 defects and DXd resistance were also verified in diverse PCa cell lines (**Fig. 3C**). These results indicate that the payload, DXd contributes to the DS-7300a resistance in TP53-deficient tumors.

**Figure 3.**
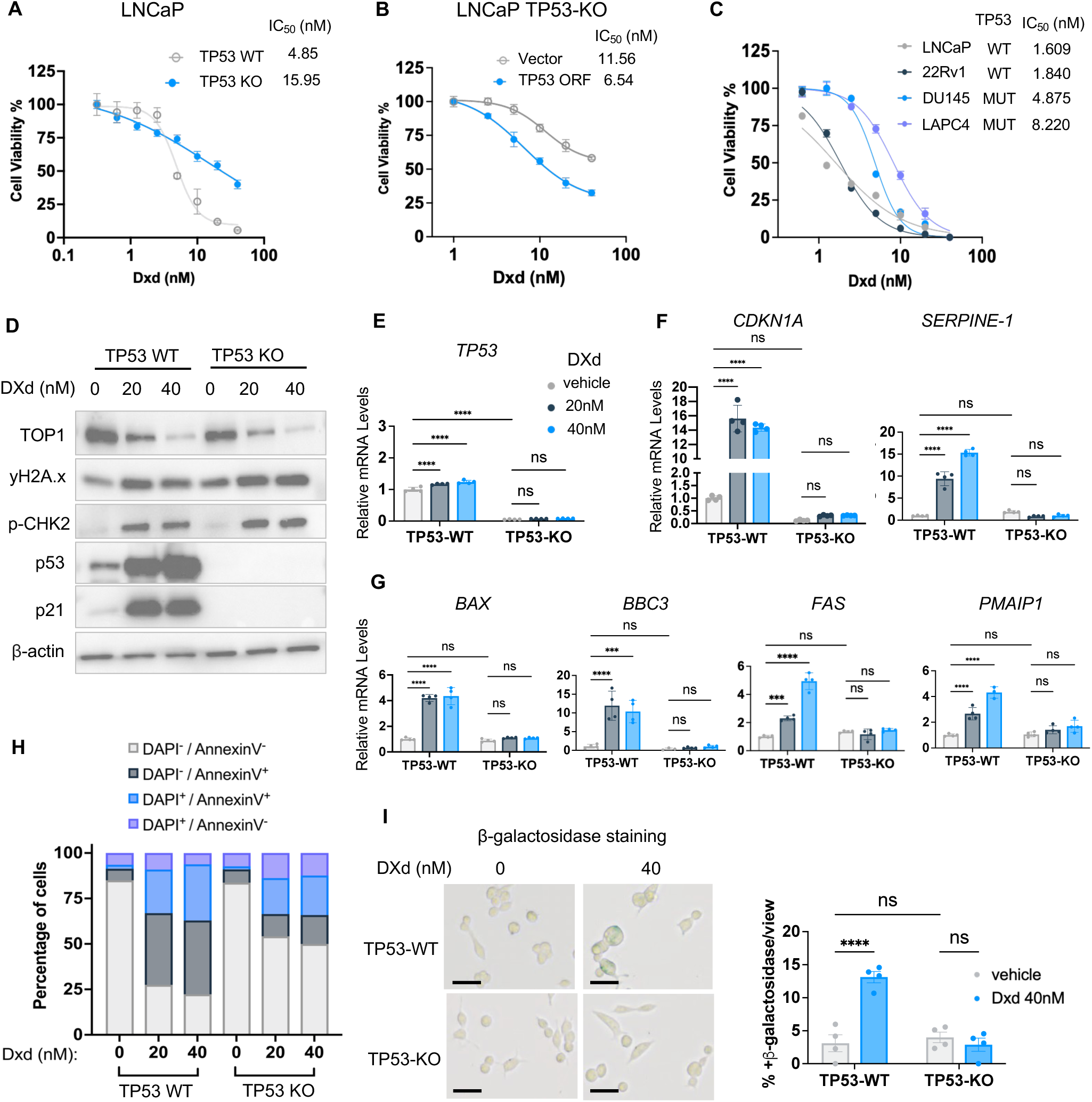
DS-7300a’s payload DXd triggers apoptosis and senescence via activating p53. **A.** Dose-response curves and IC_50_ of DXd in LNCaP with or without TP53 KO. **B.** Re-introducing TP53 ORF sensitizes TP53-KO LNCaP cells to DXd. **C.** Comparison of DXd IC_50_ in various PCa cell lines containing wildtype or mutated *TP53.* **D.** Western blot determining the expression of p53, p21, and DNA damage response signaling in TP53-WT and TP53-KO cells treated with DXd. **E-G.** mRNA expression levels of TP53 (E) and its direct target genes involved in senescence (F) and apoptosis (G) in TP53-WT and TP53-KO cells after DXd treatment, determined by qPCR. **H-I.** TP53-WT and TP53-KO LNCaP cells were treated with DXd at indicated concentrations, followed by Annexin V and DAPI analysis using Flow (H) or beta-galactosidase staining (I). Scale bar = 50um. Cells were treated with DXd for 48 hours in all experiments. Data represent the mean ± standard deviation of triplicates. IC_50_ values were calculated using GraphPad Prism version 10.4.1. *P* values determined by one-way ANOVA analysis; *P < 0.05; **P < 0.01; ***P < 0.001; ****P < 0.0001; ns, not significant.

TOP1 inhibitors, such as camptothecin derivatives, form irreversible TOP1-cleavable complexes (TOP1cc), cause DNA strand breaks, and trigger the DNA damage response [13, 38]. As camptothecin derivatives, DXd and DXd-based ADCs have been shown to induce DNA damage in cancer cells [14, 16–18]. We observed that DXd treatment led to the degradation of the TOP1 protein and the activation of DNA damage response signaling, characterized by CHK2 phosphorylation and γH2AX formation (**Fig. 3D**). DXd dramatically elevates p53 at the protein level but has a limited impact on its mRNA level (**Fig. 3D, E**), indicating a role of DXd in stabilizes p53 protein. This finding is consistent with prior reports that the ATM/ATR/CHK pathway triggers p53 phosphorylation and tempers its interaction with MDM2, leading to p53 protein stabilization and activation [39–41]. In addition, p53 transcriptional activity was enhanced after DXd treatment, as featured by the induction of p53 direct target genes that are involved in apoptosis (NOXA/*Pmaip1*, PUMA/*Bbc3*, BAX, and FAS) and cellular senescence (p21/*CDKN1A* and PAI-1/*Serpine1*) (**Fig. 3F-G)**.

In addition, we compared the effects of DXd in TP53-wildtype and knockout cancer cells. Although DXd exhibited similar effects on the CHK2 activation and DNA damage in both cells (**Fig. 3D**), it failed to induce the expression of apoptosis and senescence markers in the TP53-deficient context (**Fig. 3D-G**), suggesting p53 serves as a downstream sensor of DXd-caused DNA damage. Along the same line, apoptosis assays using Annexin V and DAPI staining revealed that, compared to TP53-wildtype cells, DXd treatment caused much fewer early apoptosis (Annexin V^+^/DAPI^−^) and late apoptosis (Annexin V^+^/DAPI^+^) in TP53-deficient cancer cells (**Fig. 3H**). Furthermore, DXd led to cellular senescence in TP53-wildtype LNCaP cells but had limited effects on TP53-deficient counterparts (**Fig. 3I**). Collectively, these results demonstrate that the TOP1 inhibitor DXd stabilizes and activates p53, which plays a vital role in triggering subsequent apoptosis and cellular senescence.

### Loss of TP53 impaired the senescence and apoptosis induced by DS-7300a

Next, we treated TP53 wild-type and knockout LNCaP cells with the B7-H3-targeting DS-7300a and assessed its effects on DNA damage response, apoptosis, and senescence. Phenocopying its payload DXd, DS-7300a treatment inhibited TOP1, increased CHK2 phosphorylation, and induced γH2AX formation regardless of TP53 status (**Fig. 4A**). DS-7300a remarkably stabilized p53 protein and activated p53 target genes involved in apoptosis and cellular senescence in a p53-dependent manner (**Fig. 4A-D**). Loss of TP53 impaired the apoptosis induced by DS-7300a, characterized by reduced BAX and FAS expression and fewer Annexin V^+^/DAPI^−^ cells (**Fig. 4D, E**). Also, DS-7300a-induced p21 upregulation and cellular senescence were abolished entirely in TP53-deficient cancer cells (**Fig. 4A, F**).

**Figure 4.**
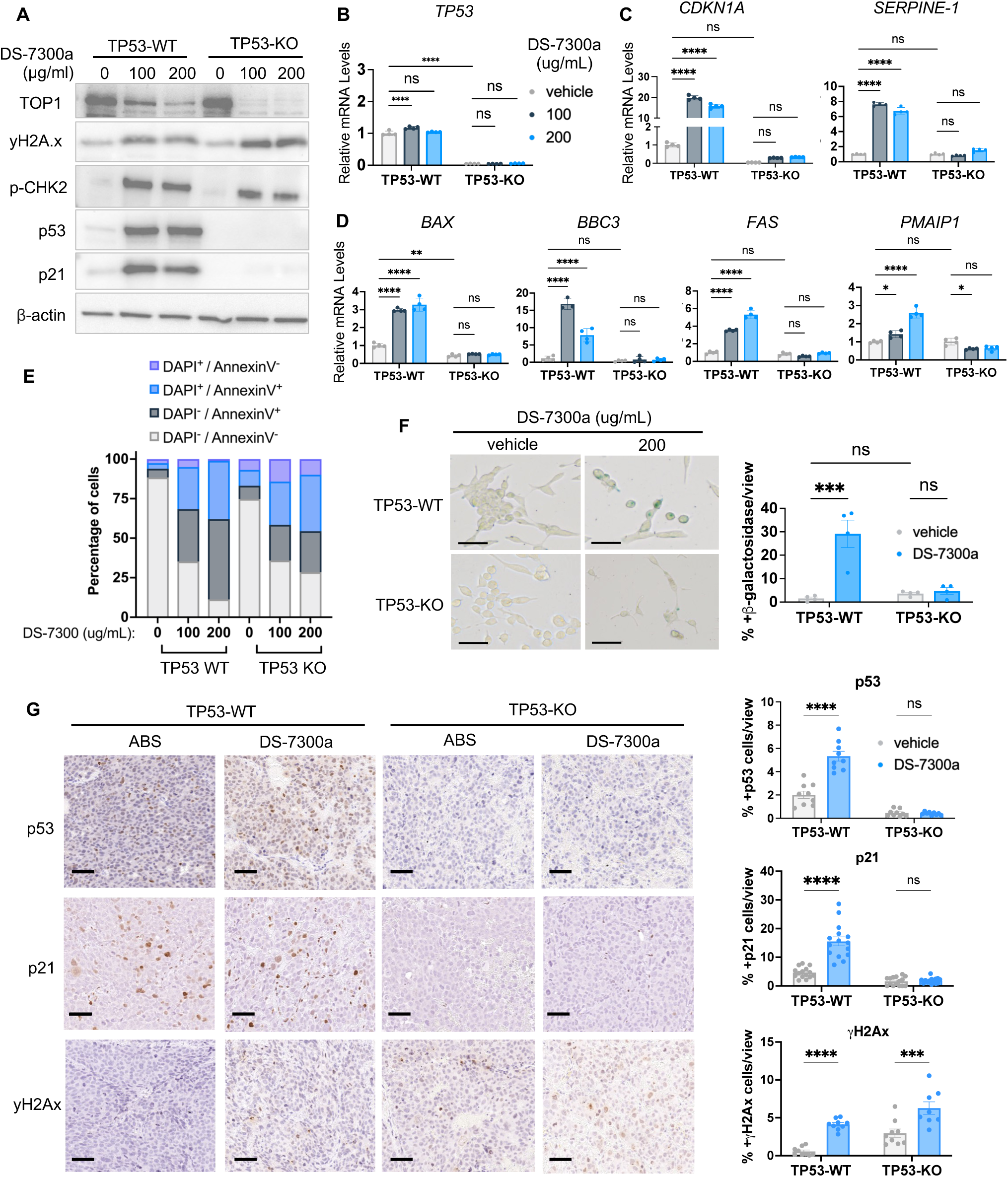
Loss of TP53 impaired the senescence and apoptosis induced by DS-7300a. **A.** Western blot determining the expression of p53, p21, and DNA damage response signaling in TP53-WT and TP53-KO cells treated with DS-7300a for 96h. **B-D.** mRNA expression levels of TP53 (B) and its direct target genes involved in senescence (C) and apoptosis (D) in TP53-WT and TP53-KO cells after DS-7300a treatment for 96h, determined by qPCR. **E-F.** TP53-WT and TP53-KO cells were treated with DS-7300a for 72h, followed by apoptosis analysis (E) or beta-galactosidase staining (F). Scale bar = 50um. **G**. Representative images and quantification of IHC staining of indicated markers in TP53-WT and TP53-KO LNCaP xenograft tumors treated with 0.5 mg/kg DS-7300a. Scale bar = 50um. Data represent the mean ± standard deviation of triplicates. Statistics were calculated using GraphPad Prism version 10.4.1. *P* values determined by one-way ANOVA analysis; *P < 0.05; **P < 0.01; ***P < 0.001; ****P < 0.0001; ns, not significant.

Furthermore, IHC staining in xenograft tumors revealed that DS-7300a treatment *in vivo* triggered DNA damage, stabilized p53 protein, and induced senescence in TP53-intact prostate tumors (**Fig. 4G**). By contrast, it had much less effect on that of TP53-deficient tumors (**Fig. 4G**). These studies uncover the molecular basis of DS-7300a’s anti-tumor activity and underscore p53’s essential role in this process. More importantly, these studies provide mechanistic insights into DS-7300a resistance in TP53-deficient tumors and guide the design of novel combinatorial strategies to improve DS-7300a’s efficacy in this context.

### Ferroptosis inducer resensitizes TP53-deficient cancer cells to DS-7300a

Ferroptosis is an iron-dependent form of regulated cell death triggered by lipid peroxidation, which is mechanistically and morphologically distinct from apoptosis [42, 43]. Glutathione peroxidase 4 (GPX4) is an antioxidant enzyme that uses glutathione (GSH) to detoxify lipid peroxidation and plays a vital role in anti-ferroptosis [43, 44]. Inhibition of GPX4 using small molecule compound RSL3 or ML162 causes the accumulation of lipid peroxidation and oxidative damage, thereby inducing ferroptosis [43].

Interestingly, DS-7300a treatment induced GPX4 expression in TP53-deficient PCa *in vitro* and *in vivo* (**Fig. 5A-B and S2B-C**). On the other hand, we found that DS-7300a treatment caused a remarkable elevation of lipid peroxidation and synergized with GPX4 inhibition to trigger lipid peroxidation, effects that could be abolished by the ferroptosis inhibitor ferrostatin (**Fig. 5C**). This finding suggests that the increased GPX4 expression might be a therapeutic vulnerability to overcome the resistance to DS-7300a in TP53-deficient cells. It prompted us to explore the combination of the ferroptosis inhibitor RSL3 and DS-7300a to counteract resistance associated with TP53 loss.

**Figure 5.**
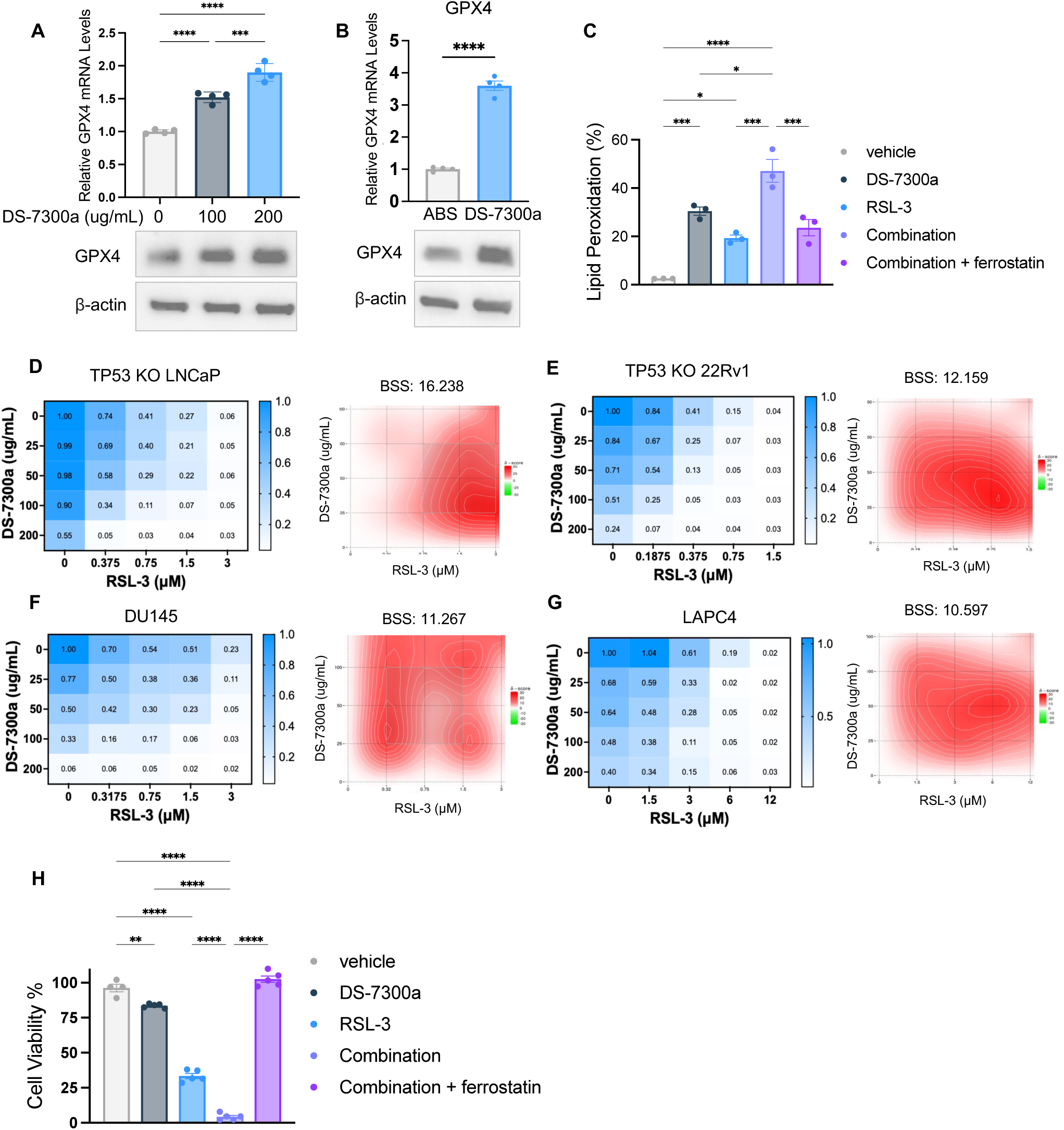
Ferroptosis inducer enhances the anti-tumor effects of DS-7300a in TP53-deficient tumors. **A-B.** GPX4 expression in TP53-KO LNCaP cells treated with DS-7300a *in vitro* (A) and TP53-KO LNCaP xenograft tumors after DS-7300a treatment *in vivo* (B), determined by Western blot and qPCR. **C.** Lipid peroxidation levels in TP53-KO LNCaP cells treated with DS-7300a or/and RSL-3 with or without ferroptosis inhibitor ferrostatin. **D-G.** Dose-response matrix (left) and drug interaction plots (right) revealing the synergistic effects of DS-7300a and RSL-3 in various *TP53-*deficient PCa cell lines: TP53-KO LNCaP (D), TP53-KO 22Rv1 (E), DU145 (TP53-mutated, F), and LAPC4 (TP53-mutated, G). Drug interaction landscapes and synergy scores were generated using SynergyFinder. **H.** Cell viability assay of TP53-KO LNCaP cells treated with DS-7300a or/and RSL-3 with or without ferroptosis inhibitor ferrostatin. Data represent the mean ± standard deviation of quadruplicates. Data represent the mean ± standard deviation of triplicates. Statistics were calculated using GraphPad Prism version 10.4.1. *P* values determined by unpaired two-tailed t test (B) or one-way ANOVA analysis (A, C, H). *P < 0.05; **P < 0.01; ***P < 0.001; ****P < 0.0001; ns, not significant.

Hence, we tested this combination in LNCaP cells with TP53 knockout. As shown in **Fig. 5D**, RSL3 exhibited strong synergistic effects with DS-7300a in TP53-deficient LNCaP cells with a Bliss synergy score of 16.2. Similar synergistic effects were also observed in 22Rv1 cells with TP53 deletion (**Fig. 5E**). Furthermore, we determined the efficacy of DS-7300a combined with RSL3 in DU145 and LAPC4 that contain TP53 mutations (**Fig. 5F-G**). These results indicated that RSL3 synergized with DS-7300a in TP53-deficient cancer cells and resensitized these cells to DS-7300a treatment. Notably, these cytotoxic effects were entirely abrogated by the ferroptosis inhibitor ferrostatin (**Fig. 5H**), suggesting that ferroptosis contributes to the cell death induced by the combination.

### Ferroptosis inducer enhances the anti-tumor effects of DS-7300a in TP53-deficient tumors

Finally, we tested the combination of DS-7300a and the ferroptosis inducer JKE-1674 in TP53-deficient humanized hB7-H3-DX1 PCa model (**Fig. 6A**). Compared to the single treatment of DS-7300a or JKE-1674, the combination effectively suppressed the progression of hB7-H3-DX1 tumors *in vivo* with limited toxicity (**Fig. 6B-D**). In addition, we tested the combination in the TP53 knockout LNCaP xenograft model (**Fig. 6A**). After tumors reached 50-70mm^3^, tumor-bearing mice were treated with ABS vehicle control or the combination of 3 mg/kg DS-7300a and 15 mg/kg JKE-1674 (**Fig. 6E**). The combination treatment remarkably attenuated the growth of TP53-deficient prostate tumors with little toxicity (**Fig. 6F-G and S2D**). Immunohistochemistry analysis revealed that the combination induced DNA damage and promoted lipid peroxidation and ferroptosis, as evidenced by elevated 4HNE levels (**Fig. 6H**). Collectively, these preclinical studies demonstrate that inducing ferroptosis can overcome resistance to DS-7300a in TP53-deficient tumors, and that combining a ferroptosis inducer with DS-7300a is more effective for treating cancers harboring TP53 defects.

**Figure 6.**
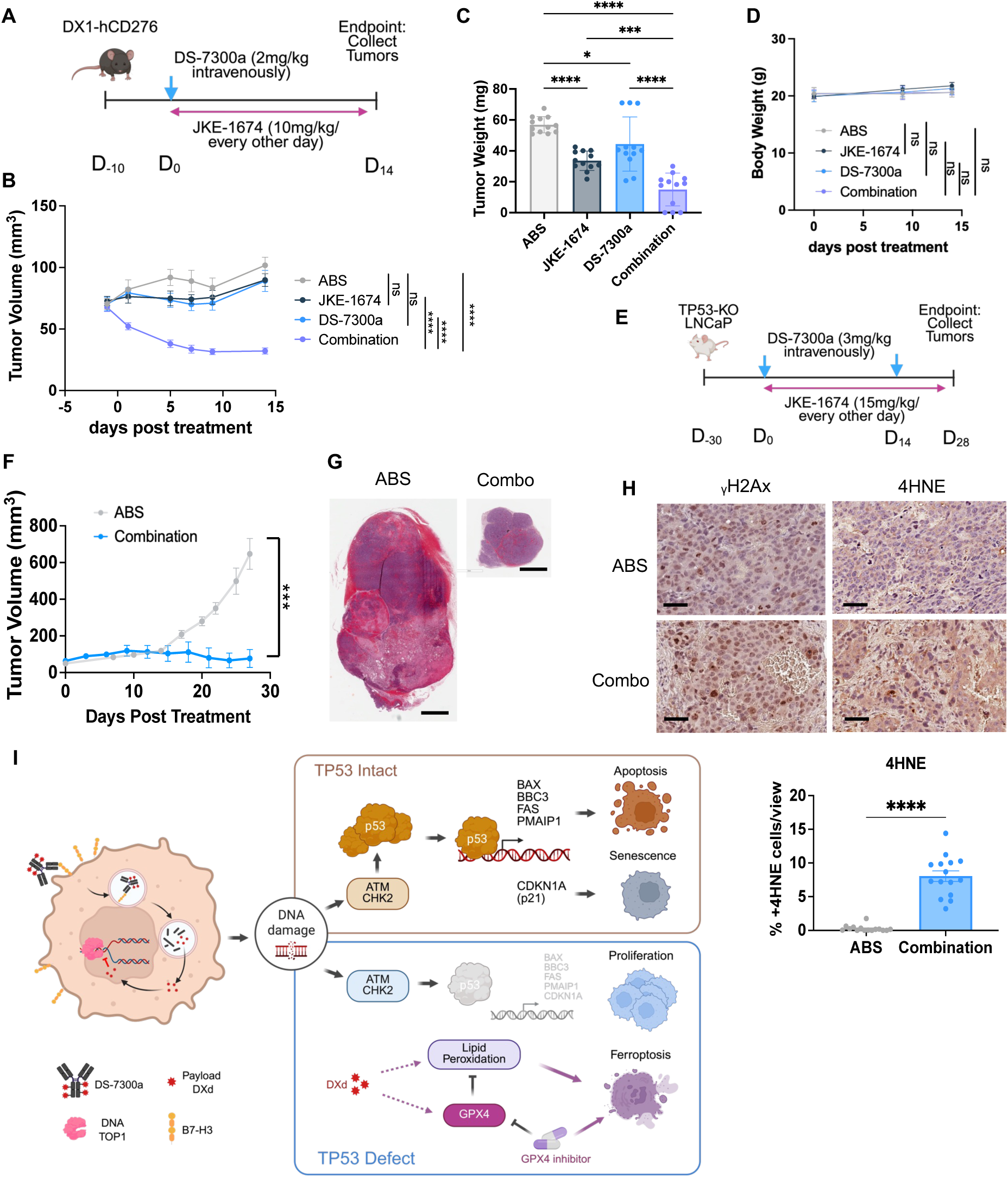
Combining DS-7300a with ferroptosis inducer suppresses TP53-deficient tumors. **A.** Schematic of experimental design of combination treatment in humanized B7-H3 models. 2×10^6^ DX1-hCD276 cells were subcutaneously injected into both flanks of male B-hB7-H3 mice, followed by treatment of single agents or combination of DS-7300a (2 mg/kg; i.v.; once) or/and JKE-1674 (10 mg/kg; oral gavage, every two days). **B-D.** Tumor growth over time (B), tumor weights at the endpoint (C), and mice body weight changes (D) of humanized B7-H3 PCa models after treatments. **E.** Schematic of experimental design in xenograft models. 2×10^6^ TP53-KO LNCaP cells were subcutaneously injected into both flanks of SCID mice. When tumors reached approximately 75mm^3^, tumor-bearing mice were randomized for combination treatment (3 mg/kg DS-7300a, i.v., biweekly, twice; 15 mg/kg JKE-1674, oral gavage, every two days) or ABS vehicle control. **F-G.** Tumor growth over time (F) and H&E staining at the endpoint (G) in xenograft models are shown. Scale bar = 2 mm. **H.** Representative images and quantification of 4-HNE IHC staining in xenograft tumors after treatments. Scale bar = 50um **I.** Schematic working model. Data represent the mean ± standard deviation of triplicates or otherwise stated. *P* values determined by unpaired two-tailed t test (F, H) or one-way ANOVA analysis (B, C, D). *P < 0.05; **P < 0.01; ***P < 0.001; ****P < 0.0001; ns, not significant.

## DISCUSSION

Our studies demonstrate that the anti-tumor activity of DS-7300a, an advanced B7-H3-targeting ADC drugs, highly depends on functional p53 in cancer cells, whereas TP53 defects confer resistance to DS-7300a (**Fig. 6I**). Mechanistically, the TOP1 inhibitor payload DXd causes DNA strand breaks and triggers DNA damage response by activating ATM/ATR/CHK signaling. In TP53-intact cells, this DNA damage stabilizes p53 and activates its downstream transcriptional program, leading to apoptosis and senescence in cancer cells. In contrast, TP53-deficient cancer cells fail to sense DXd-induced DNA damage and maintain a high proliferation rate, lowering their sensitivity to DXd and DS-7300a (**Fig. 6I**). Furthermore, we reported a novel effect of DS-7300a treatment, elevating lipid peroxidation and the antioxidant enzyme GPX4 in TP53-deficient cancer cells, as an actionable therapeutic vulnerability. More importantly, our preclinical studies in isogeneic and humanized B7-H3 PCa models demonstrated that pharmacological inhibition of GPX4 potentiates the anti-tumor effects of DS-7300a in *TP53*-deficient tumors by inducing ferroptosis (**Fig. 6I**).

The immune checkpoint B7-H3 has emerged as a therapeutic target across diverse cancer types. Enoblituzumab (MGA271), a mAb against B7-H3, is currently under phase I/II clinical investigation for refractory solid tumors (NCT01391143, NCT02923180, NCT02982941, NCT02381314, NCT02475213, NCT06014255, and NCT04630769) [45, 46]. Our recent study in preclinical models found that loss of PTEN and/or TP53 induces B7-H3 expression in cancer cells, and that inhibiting B7-H3 with mAb exerts anti-tumor effects in PTEN/TP53-deficient PCa by enhancing intratumoral infiltration of T cells and NK cells [36]. Considering DS-7300a’s anti-tumor efficacy correlates with higher B7-H3 expression in cancer cells in preclinical models [16, 37], initially, we hypothesized that TP53-deficient tumors were more sensitive to DS-7300a due to the elevated B7-H3 expression. Surprisingly, our functional and mechanistic studies demonstrate that TP53 defects confer resistance to DS-7300a, and p53 plays an essential role in mediating the cytotoxic activity of DS-7300a and its payload DXd. The distinct effects of TP53 defects on the efficacy of B7-H3-targeting mAb versus ADC underscore the need for caution when identifying molecular biomarkers to predict response to different types of drugs targeting the same protein. Elucidating the mechanism of action of individual drugs is equally vital as evaluating their anti-tumor activities in preclinical and clinical studies.

As noted above, DS-7300a has been clinically tested in different types of refractory cancers. It is crucial to identify patient subsets that are most likely to benefit from DS-7300a treatment. TP53 is one of the most frequently altered tumor suppressor genes in human cancers. Our studies, report TP53 as a genetic determinant that impacts the response to DS-7300a. As a multi-institutional DS-7300a clinical study has recently been initiated in advanced PCa patients (NCT06863272), our studies provide novel and timely insights into using TP53 status as a potent predictive marker for DS-7300a treatment in PCa among other cancer types. Future studies are warranted to assess the association between TP53 status and DS-7300a response in a clinical setting. If successful, it may inform patient selection for future clinical trials of DS-7300a and facilitate biomarker identification for its clinical use.

TP53 mutation or loss leads to a more aggressive tumor phenotype and affects the efficacy of conventional therapies, including radiotherapy, chemotherapy, and hormone therapy [47, 48]. Our studies demonstrated that TP53 defects may contribute to resistance to ADC, a new class of anti-tumor drugs that combine chemotherapy with immunotherapy. Notably, the DS-7300a resistance in TP53-deficient tumors is attributed to the DNA-damaging payload (TOP1 inhibitor, DXd) rather than the target B7-H3. It suggests that, in addition to DS-7300a, TP53 defects might confer resistance to other DXd-based ADCs. However, it may or may not apply to ADCs with other payload types, such as microtubule-targeting agents or RNA transcription inhibitors. Future studies are needed to test these hypotheses and determine if they are more efficient for treating TP53-deficient tumors. Besides, radionuclide antibody conjugate (RAC), combining antibodies and radionuclide payload, has become an emerging tool for cancer diagnosis and therapeutics [49, 50]. Given the nature of radionuclides in inducing DNA damage, it would be interesting to determine TP53’s impact on the efficacy of RAC in preclinical models.

Distinct from apoptosis, ferroptosis is a newly discovered form of cell death triggered by lipid peroxidation [42, 43]. Ferroptosis has sparked great interest in the cancer research community because inducing ferroptosis holds great potential for treating cancers refractory to conventional therapies. Prior studies have shown that ferroptosis inducers can restore the sensitivity of radiotherapy- and chemotherapy-resistant tumors to these agents [51–54]. By illuminating the mechanism underlying DS-7300a resistance in TP53-deficient tumors, we developed a novel approach to enhance DS-7300a efficacy by inducing ferroptosis. Our preclinical studies provide “proof-of-concept” evidence supporting ferroptosis as a therapeutic vulnerability in ADC-resistant cancers and the therapeutic potential of combining ferroptosis inducers with other DXd-based ADC in TP53-deficient cancers. Future studies are warranted to elucidate the mechanisms by which DS-7300a reprograms lipid metabolism and the anti-ferroptosis system in TP53-deficient cancer cells. It is worth noting that ferroptosis also contributes to p53-mediated radiosensitization, and ferroptosis inducers render p53-mutant tumors more sensitive to radiotherapy [54]. Therefore, combining ferroptosis inducers might be a universal strategy to overcome the resistance to DNA-damaging agents in malignancies with TP53 deficiency.

In summary, our studies demonstrate that p53 status dictates anti-tumor responses to DS-7300a, and ferroptosis induction represents a promising therapeutic approach to enhance the efficacy of DXd-based ADCs in malignancies harboring TP53 defects. Our studies may accelerate the development of effective therapies for PCa patients with TP53 defects and provide a strong rationale for using TP53 status as a molecular biomarker to guide patient selection for clinical application of DS-7300a. This biomarker-driven combination strategy may also yield novel insights into the clinical trial design of other DXd-based ADCs and ferroptosis inducers in advanced PCa and other malignancies.

## Reporting Summary

Further information on research design is available in the Nature Portfolio Reporting Summary linked to this article.

## Data Availability

Source data for Western Blot images and Numerical results in all Figures and Extended Data Figures will be provided as Source Data files. All other data supporting the findings of this study are available from the corresponding author upon reasonable request.

## Authors’ Contributions

J.L. and D.Z. designed experiments, analyzed results, created figures, and wrote the manuscript. J.L. performed the experiments. F.C assisted with animal studies, synergism assays, and technical support. W.S. generated the TP53 knockout isogenic cell lines and provided technical support. X.L. provided mouse husbandry and assisted with animal experiments. C.M., Q.G., Y.A., and Z.F provided technical support. J.Z. helped with processing histopathological samples. B.G., S.G., K.C., D.F., and A.A. provided intellectual contributions throughout the project.

## Acknowledgments

We thank Daiichi Sankyo Inc. for providing DS-7300a and ABS vehicle control for this study. We acknowledge the MD Anderson Small Animal Imaging Facility (supported by NIH/NCI P30 CA016672 Cancer Center Support Grant). Y.A. is an MD Anderson CATALYST summer trainee, supported by a CPRIT Research Training Award (RP210028). B. Gan has been supported by RP230072 from the Cancer Prevention & Research Institute of Texas (CPRIT), and R01CA181196, R01CA244144, R01CA247992, R01CA269646, and U54CA274220 from the National Institutes of Health. D. Zhao is a CPRIT Scholar in Cancer Research and has been supported by the CPRIT Recruitment of First-Time Tenure-Track Faculty Award RR190021, NIH/NCI R01 CA275990 and CA278889, Prostate Cancer Foundation Challenge Award FP00016492, and DoD CDMRP IDA award PC230358.

