## Supplementary Figures 1-2 for "Improve the Efficacy of B7-H3-Targeting Antibody-Drug Conjugate DS-7300a in TP53-deficient Tumors by Inducing Ferroptosis"

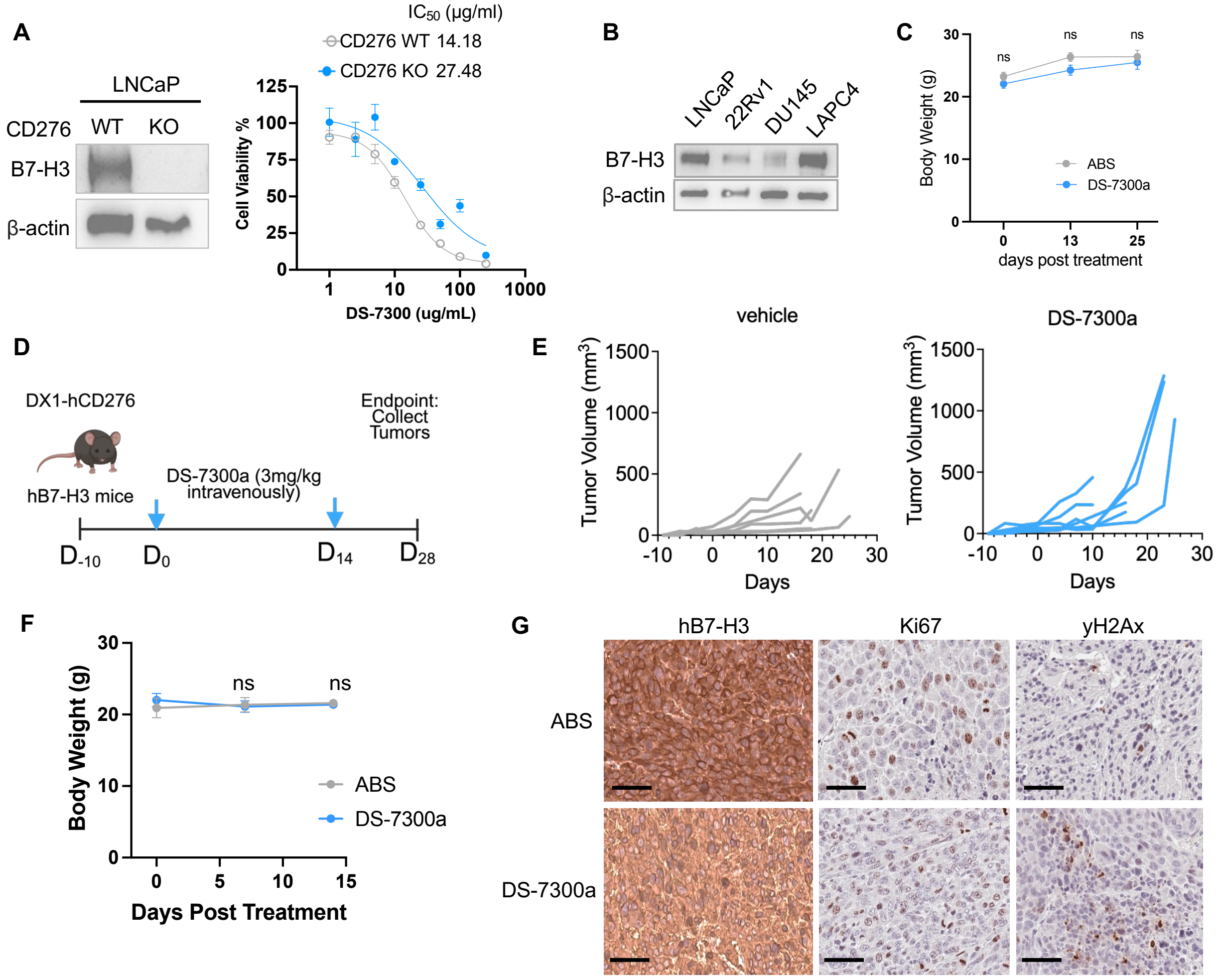


**Supplementary Figure 1. TP53 defects confer resistance to DS-7300a, a B7-H3-targeting ADC.**

**A**. Western blot of B7-H3 and dose-response assays present the sensitivity of CD276-wildtype (WT) and CD276-knockout (KO) LNCaP cells to DS-7300a. $IC_{50}$ values are shown **B.** Western blot analysis of B7-H3 protein in various PCa cell lines containing various TP53 status. **C**. Images of TP53-WT and TP53-KO LNCaP xenograft tumors after ABS or DS-7300a treatment (0.5 mg/kg) and body weight of tumor-bearing mice treated with ABS or DS-7300a (0.5 mg/kg) over the course of the study. **D.** Schematic of experimental design using humanized B7-H3 models. $2\times10^6$ DX1-hCD276 cells were subcutaneously injected into both flanks of male B-hB7-H3 mice, followed by treatment of ABS vehicle control or DS-7300a (3 mg/kg; i.v., biweekly, twice). **E-F.** Tumor growth curves of individual DX1-hCD276 tumors (E) and body weights of mice (F) treated with ABS or DS-7300a (3 mg/kg). **G.** Immunohistochemistry analysis of hB7H3, Ki67 and DNA damage markers in DX1-hCD276 tumors after treatment. Scale bar = 50um. Data represent the mean ± standard deviation of triplicates or otherwise stated. *P* values determined by unpaired two-tailed t test; *P < 0.05; **P < 0.01; ***P < 0.001; ****P < 0.0001; ns, not significant.

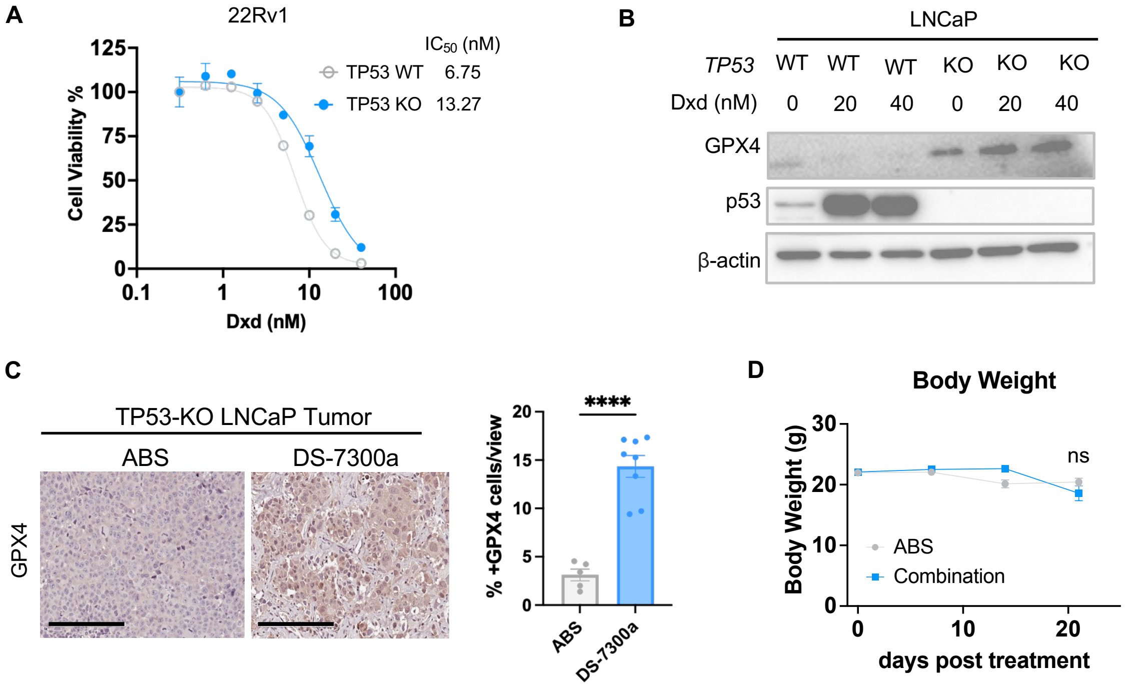


**Supplementary Figure 2. Ferroptosis inducer enhances the anti-tumor effects of DS-7300a in TP53-deficient tumors.**

**A**. Dose-response curves and $IC_{50}$ of DXd in 22Rv1 with or without TP53 KO. **B.** Western blot analysis of GPX4 and p53 in TP53-WT and TP53-KO LNCaP cells after DXd treatment. **C.** Representative images and quantification of GPX4 IHC staining in TP53-KO LNCaP tumors after DS-7300a treatment. Scale bar = 200um. **D.** Body weight of mice treated with ABS or combination of DS-7300a and JKE-1674 in TP53-KO LNCaP xenograft model. Data represent the mean ± standard deviation of triplicates or otherwise stated. *P* values determined by unpaired two-tailed t test; *P < 0.05; **P < 0.01; ***P < 0.001; ****P < 0.0001; ns, not significant.
